# Human neutrophils direct epithelial cell extrusion to enhance intestinal epithelial host defense during *Salmonella* infection

**DOI:** 10.1101/2022.03.25.485742

**Authors:** Anna-Lisa E. Lawrence, Ryan P. Berger, David R. Hill, Sha Huang, Veda K. Yadagiri, Brooke Bons, Courtney Fields, Gautam J. Sule, Jason S. Knight, Christiane E. Wobus, Jason R. Spence, Vincent B. Young, Mary X. O’Riordan, Basel H. Abuaita

## Abstract

Infection of the human gut by *Salmonella enterica* Typhimurium (STM) results in a localized inflammatory disease that is not mimicked in murine infections. To determine mechanisms by which neutrophils, as early responders to bacterial challenge, direct inflammatory programming of human intestinal epithelium, we established a multi-component human intestinal organoid (HIO) model of STM infection. HIOs were micro-injected with STM and then seeded with primary human polymorphonuclear leukocytes (PMN-HIOs), specifically neutrophils and analyzed for bacterial growth and host cell survival. Surprisingly, PMNs did not affect luminal colonization of *Salmonella,* but their presence reduced intraepithelial bacterial burden. Adding PMNs to infected HIOs resulted in substantial accumulation of shed intestinal epithelial cells that could be blocked by Caspase-1 or Caspase-3 inhibition. Cleaved Caspase-3 was present in epithelial cells, but expression of the inflammasome adaptor, ASC, was only detected in PMNs. Caspase inhibition also increased bacterial burden in the epithelium of the PMN-HIO, suggesting PMNs enhance activation of cell death pathways in human intestinal epithelial cells as a protective response to infection. These data support a critical function for neutrophils beyond their antimicrobial role whereby they amplify cell death and extrusion of epithelial cells from the *Salmonella*-infected intestinal monolayer.

**Significance statement:** Neutrophils are early responders to *Salmonella* intestinal infection, but how they influence infection progression and outcome is unknown. Here we use a co-culture model of human intestinal organoids and human primary neutrophils to study the contribution of human neutrophils to *Salmonella* infection of the intestinal epithelium. We found that neutrophils markedly enhanced epithelial defenses, including enhancing cell extrusion to reduce intraepithelial burden of *Salmonella* and association with the epithelium, rather than directly killing *Salmonella* in the HIO lumen. These findings reveal a novel role for neutrophils in the gut beyond killing invading pathogens and illuminate how neutrophils can reprogram cells in the gut environment to enhance antimicrobial defenses.

## Introduction

*Salmonella enterica* is one of the most common causes of foodborne disease, responsible for an estimated 1.35 million infections in the United States each year (1). *S. enterica* serovar Typhimurium (STM), one of the most prevalent *S. enterica* serovars, infects via the fecal-oral route and stimulates robust inflammation in the host intestinal milieu, leading to gastroenteritis and diarrheal disease (2). *Salmonella* pathogenesis is commonly studied *in vivo* using mouse infection models and now more recently *in vitro* using human derived organoid and enteroid models (3–7). Through these studies a critical role for epithelial cell death and extrusion of *S. enterica* infected cells has been shown to regulate infection outcome by reducing epithelial bacterial burden and restricting the infection to the intestine (5, 8–11).

Epithelium intrinsic induction of cell death and extrusion pathways are essential in maintenance of normal intestinal homeostasis during health and infection, however it is also known that innate immune cells play a dominant role in determining infection outcome by bacterial infections, including *Salmonella* (12, 13). One of the earliest responders and the most abundant cell types found in *Salmonella*-infected individuals are polymorphonuclear leukocytes (PMNs), specifically neutrophils (14, 15). PMNs defend against bacterial infections through multiple mechanisms: antimicrobial effectors like degradative proteases and ion chelators, production of reactive oxygen species and formation of sticky antimicrobial neutrophil extracellular traps (NETs) (16). And while PMNs are very effective at killing extracellular bacteria (17), the role of PMNs in a more complex infection where a large proportion of bacteria reside within host cells, like the intestinal epithelium, is unknown.

More recently, it is becoming appreciated that PMNs serve other functions beyond killing extracellular pathogens, e.g., changing the microenvironment via molecular oxygen depletion, regulating nutrient availability, and through production of inflammatory mediators (18, 19). Notably, Gopinath et al. found that neutrophilia induced a super-shedder phenotype in a mouse infection model of *Salmonella* (20), but how the interaction between epithelial cells and PMNs affects the outcome of bacterial infections is still poorly understood. To address this gap in knowledge, we generated a co-culture model of primary human PMNs, specifically neutrophils, with human intestinal organoids (HIOs) termed PMN-HIOs to study the contribution of PMNs during infection with *Salmonella enterica* serovar Typhimurium (STM).

Using this PMN-HIO model, we evaluated how PMNs modulate intestinal epithelial host defenses during infection, compared to infected HIOs alone. We show here that the presence of PMNs elevates the overall inflammatory tone of the epithelium and markedly promotes cell death and extrusion of epithelial cells, thereby reducing *Salmonella* intraepithelial burden.

## Results

### Human PMNs transmigrate into the HIO lumen during infection and reduce *Salmonella* intraepithelial burden

PMNs are known to transmigrate across intestinal epithelial layers during early stages of inflammation (21, 22), therefore we asked whether PMNs would transmigrate into the HIO lumen during infection. 10^5^ *Salmonella enterica* Typhimurium (STM) were microinjected into the HIO lumen and cultured with PMNs isolated from healthy human volunteers for 8h (PMN-HIOs). To quantify PMN recruitment to infected HIOs, PMNs were pre-labeled with Carboxyfluorescein succinimidyl ester (CFSE) prior to co-culture with HIOs. PMN-HIOs were collected at 8h post-infection (hpi), washed to remove unassociated neutrophils, dissociated into a single cell suspension and the percentage of CFSE-positive cells was enumerated by flow cytometry. There was a significant increase in the number of PMNs associated with infected HIOs compared to PBS controls, with approximately 5% of total cells present in PMN-HIOs staining positive for CFSE (Fig. 1A). Immunofluorescent staining for neutrophil-specific Myeloperoxidase (MPO) was performed on paraffin sections to further monitor localization of PMNs within PMN-HIOs. In contrast to PBS-injected controls, MPO-positive cells were observed in the lumen of STM-infected HIOs, confirming that PMNs transmigrate into the HIO lumen during infection (Fig. 1B). Since PMNs are potent killers of bacterial pathogens, we tested whether PMNs controlled *Salmonella* colonization within the HIO. Although PMNs killed STM in pure PMN cultures, with ~30% of STM killed by 4hpi (SI Appendix, Fig. S1), PMNs did not alter the total levels of STM in the HIOs (Fig. 1C). This was not due to lack of PMN activation in the HIOs as we detected formation of NETs in the lumen of STM-infected PMN-HIOs (SI Appendix, Fig. S2). In addition, culture supernatants were analyzed for production of antimicrobial effectors via ELISA (SI Appendix, Fig. S3). Some antimicrobial effectors such as Elafin (PI3), a small cationic peptide secreted at mucosal surfaces (23), and Calprotectin (S100A8 and S100A9) were produced at higher levels in PMN-HIOs, compared to HIOs alone, confirming that there was no defect in the antimicrobial response to STM and in contrast suggests that PMNs augment epithelial host defenses. However, since *Salmonella* has evolved mechanisms to overcome Calprotectin-mediated immunity and thrive under these conditions; upregulation of these specific antimicrobial effectors is likely insufficient to reduce *Salmonella* colonization in the PMN-HIOs (24, 25).

**Fig. 1.**
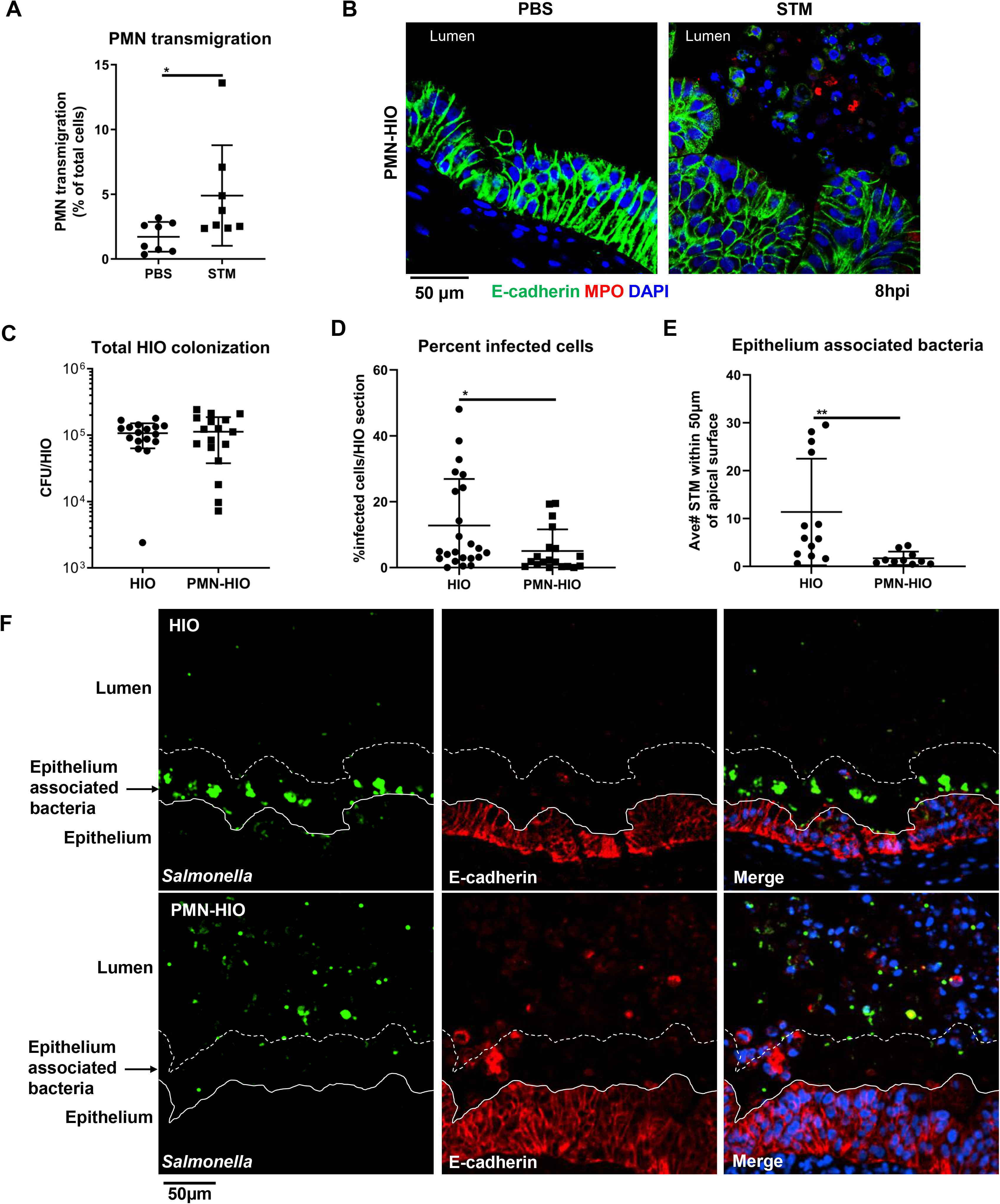
PMNs migrate into the lumen during infection and reduce both the number of infected epithelial cells and the association of bacteria with the epithelial surface. (A) Transmigration of PMNs into the lumen of HIOs was quantified using flow cytometry. HIOs were microinjected with STM or PBS and co-cultured with CFSE-labeled PMNs for 8h. PMNs-HIOs were washed to remove any unassociated PMNs, dissociated into a single cell suspension and subjected to flow cytometry. Percentage of PMNs relative to total cells acquired per PMN-HIO was determined by FlowJo software. (B) Immunofluorescent staining of HIOs microinjected with PBS or STM and co-cultured with PMNs. E-cadherin (green) marks the epithelial lining, MPO (red) is specific to PMNs and DNA is stained with DAPI. (C) Total bacterial burden per HIO or PMN-HIO was enumerated at 8hpi. (D) Quantitation of percent infected cells/HIO or PMN-HIO based on 3 fields per view per HIO. (E) Quantitation of epithelium associated bacteria. Number of bacteria within 50μm of the apical epithelial surface based on E-cadherin staining were counted and normalized per 100μm distance. (F) Representative immunofluorescent staining of HIOs and PMN-HIOs infected with STM. *Salmonella* is stained in green, E-cadherin to mark epithelial cells is shown in red, and DNA stained with DAPI in blue. Graphs show the mean and SD of n≥ 10 HIOs represented by dots from at least two independent experiments. Outliers were removed using the ROUT method with Q=0.1%. Unless otherwise stated, significance was determined by Mann-Whitney test with *p<0.05, **p<0.01.

During intestinal infection, *Salmonella* reside in both the lumen of the intestine and within epithelial cells, and it has been suggested that the intracellular pool of bacteria are important for reseeding the gut lumen to prolong infection and promote fecal shedding (8, 26). To assess what impact PMNs have on the intracellular bacterial burden, paraffin sections of *Salmonella-*infected HIOs and PMN-HIOs were stained to detect both epithelial cells and *Salmonella* and the epithelial bacterial burden was quantified by fluorescence microscopy (Fig. 1D, 1E). This analysis revealed that there were significantly fewer intracellular bacteria in the epithelial lining of PMN-HIOs compared to HIOs alone indicating that PMNs aid in reducing epithelial cell bacterial burden. Interestingly, we also observed a reduction in epithelial-surface associated bacteria suggesting that PMNs also reduce STM attachment to further protect the epithelial lining. Together, these results show that although PMNs transmigrate into the HIO lumen during *Salmonella* infection, they do not directly kill *Salmonella*, but instead enhance epithelial defenses to reduce the bacterial burden within the epithelial layer.

### PMNs enhance shedding of epithelial cells during *Salmonella* infection

In addition to measuring a significant reduction in intraepithelial bacterial burden and association of STM with epithelial cells, we observed robust accumulation of DAPI-positive cells in the lumen of STM-infected PMN-HIOs that were negative for the PMN marker MPO (Fig. 1B, F). Shedding of *Salmonella*-infected cells from the gut via programmed cell death pathways is an important defense mechanism used to protect the host from invasive *Salmonellosis* and helps reduce intestinal bacterial burden to resolve the infection and so we hypothesized that in a human infection model PMNs may enhance this process (5, 8–11). To determine whether these luminal cells were dead epithelial cells shed from the HIO epithelial lining, we performed Terminal deoxynucleotidyl transferase dUTP nick end labeling (TUNEL) on HIOs and PMN-HIOs microinjected with either STM or PBS control. The presence of PMNs induced robust accumulation of TUNEL-positive cells in the lumen of infected HIOs (Fig. 2A, 2B). While we also detected a substantial number of TUNEL-positive cells in the mesenchyme this phenotype was present in all conditions so likely was not caused by either *Salmonella* or PMNs. Accumulation of luminal TUNEL-positive cells was selectively induced by PMNs during infection, as neither infected HIOs or uninfected PMN-HIOs showed this phenotype. To confirm that these cells were epithelial cells, we stained for the epithelial marker E-cadherin, and found that the vast majority of TUNEL-positive cells in PMN-HIOs were epithelial cells (Fig. 2C, 2D). Previously we reported that STM infection in HIOs induces significant induction of TUNEL-positive cells that are retained in the epithelial lining (4), consistent with these findings, our results suggest that PMNs are enhancing shedding of these TUNEL-positive cells from the monolayer as we observe very few TUNEL-positive cells in the epithelial lining of infected PMN-HIOs (Fig. 2D).

**Fig. 2.**
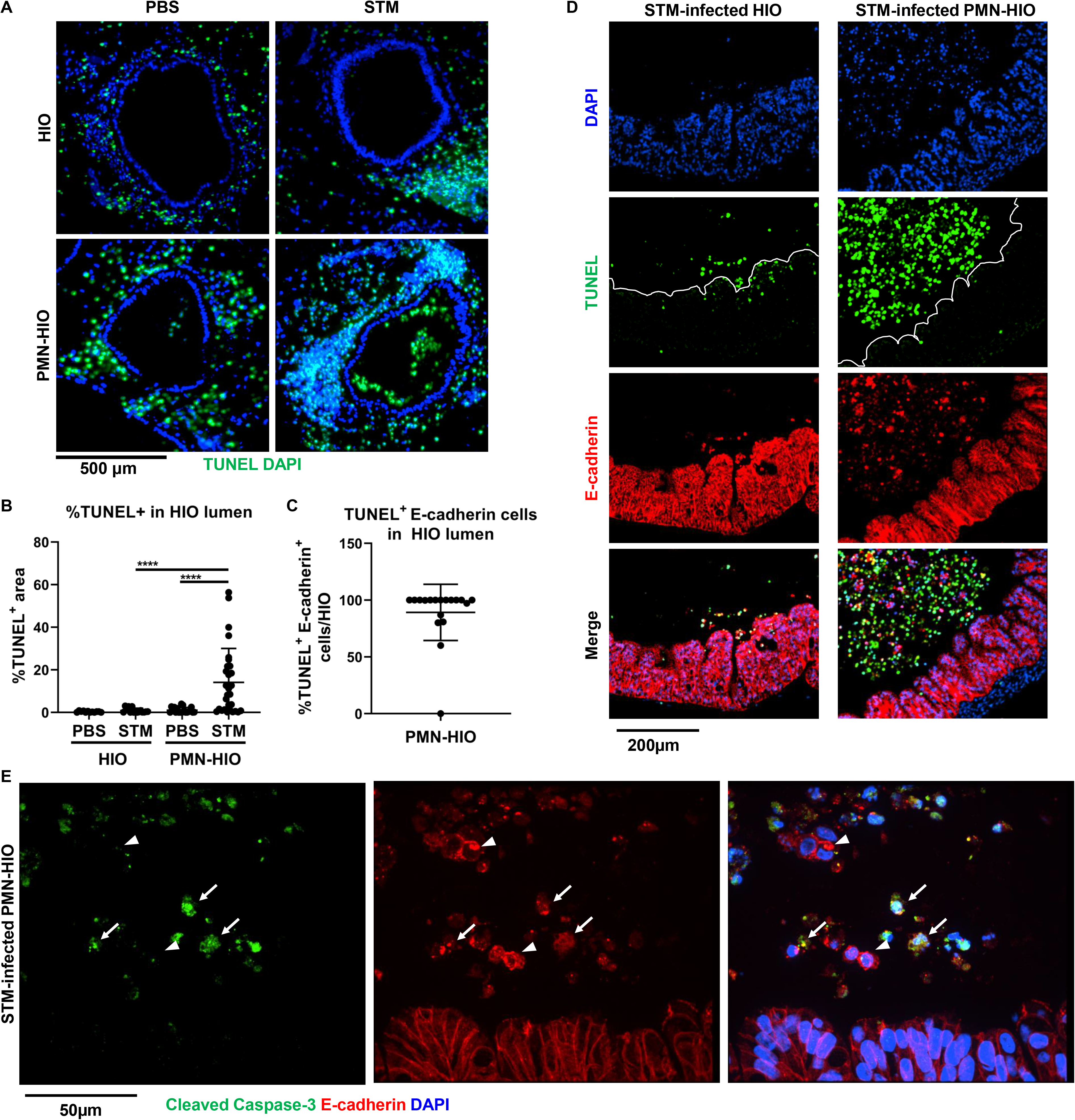
PMNs induce epithelial cell death and shedding during *Salmonella* infection. (A) Immunofluorescent images of TUNEL staining of histology sections of HIOs and PMN-HIOs injected with PBS or STM at 8hpi. (B) Quantitation of TUNEL positive cells in the lumen of HIOs and PMN-HIOs from (A). Graphs show the mean and SD of HIOs from 2 independent experiments with n>12 HIOs per group. (C) Quantitation of percent of TUNEL-positive epithelial cells in HIO lumen. The percentage of TUNEL-positive cells that stained positive for E-cadherin in the HIO lumen were assessed. (D) Representative confocal microscopy images of histology sections from STM-injected HIOs or PMN-HIOs at 8h. Sections were co-stained with TUNEL (green), epithelial cell marker E-cadherin (red), and DNA marker DAPI (blue). (E) Confocal microscopy images of histology sections of HIOs and PMN-HIOs that were stained for E-cadherin (red), cleaved Caspase-3 (green), and DNA (blue). Arrows point to cleaved Caspase-3 positive epithelial cells whereas arrowheads point to cleaved Caspase-3 negative luminal epithelial cells. Outliers were removed using the ROUT method with Q=0.1%. Significance was determined via one-way ANOVA with post-Tukey’s test for multiple comparisons where ****p<0.0001.

TUNEL staining is classically associated with apoptotic pathways, and since activated PMNs have been shown to induce apoptotic processes in lung epithelium (27), we hypothesized that PMNs induce epithelial cell apoptosis to reduce bacteria associated with the epithelial lining. To test our hypothesis, we stained STM-infected PMN-HIO sections for cleaved Caspase-3 as a marker of apoptosis. We found that many, but not all luminal epithelial cells were positive for cleaved Caspase-3 (Fig. 2E) suggesting that multiple forms of cell death were occurring in the PMN-HIO likely including Caspase-4/5 mediated shedding of *Salmonella-*infected epithelial cells as has previously been established in a human enteroid infection model (5, 6) and more recently shown in a Caco-2 model of *Salmonella* infection (28). Interestingly, PMN-induced cell shedding was not restricted to infected cells, as we observed both infected and uninfected cells in the PMN-HIO lumen (SI Appendix, Fig. S4). Together these results suggest that PMNs promote programmed cell death pathways and epithelial cell shedding during *Salmonella* infection.

### Inflammasome activation and IL-1 production is mediated by PMNs during infection

PMNs are known to affect epithelial cell function through multiple mechanisms including through NET formation. Inflammasome activation is known to occur in PMNs during bacterial infections (29) and it is known that inflammasomes also play a key role in regulating intestinal inflammation (30). While activation of noncanonical inflammasomes requiring Caspase-4/5 in human epithelial cells is a hallmark of *Salmonella* infection, activation of Caspase-1 dependent inflammasomes in innate immune cells is also critical for effective responses against invading pathogens (31). To assess how PMNs shape inflammasome activation in the PMN-HIO model, we first examined gene level expression data from RNA-seq data of HIOs and PMN-HIOs microinjected with PBS or STM (SI Appendix, Table S1). We found that PMNs significantly contributed to upregulation of several genes in inflammasome/cell death signaling during STM infection (Fig. 3A). While there was weak upregulation of IL-1β and IL-1α in STM-infected HIOs alone, we did not observe significant changes in expression of other mediators or machinery required for NLRP3 inflammasome assembly such as CASP1, NLRP3, or PYCARD (encoding ASC which was not differentially expressed under any condition) (Fig. 3A). However, consistent with previous reports studying cell death responses to STM-infection in human epithelial models, we did measure a significant increase in expression of Caspase-4 and Caspase-5. In contrast, when PMNs were added to infected HIOs, we observed stronger upregulation of *IL-1* genes and effectors involved in inflammasome activation including the upregulation of *NLRP3* and *Caspase-1* (*CASP1*). To further characterize this phenotype, we collected culture supernatants from HIOs and PMN-HIO and quantified levels of IL-1 family cytokines during infection (Fig. 3B). IL-1β or IL-1α was undetectable in infected HIOs; release of these cytokines required the presence of PMNs as IL-1β or IL-1α levels significantly increased in STM-infected PMN-HIOs. These results suggest that PMNs significantly contribute to production of IL-1 cytokines in this infection model. This is consistent with previous reports that the human epithelium is not a dominant source of IL-1 cytokines (32) and that Caspase-4/5 activation is not classically associated with significant IL-1β processing (33). Interestingly, we also observed production of IL-1RA, the antagonist of the IL-1 receptor which is usually co-expressed with IL-1α/β (34), in infected PMN-HIOs revealing an additional role for PMNs in inducing signaling processes that tune the magnitude of immune activation. In contrast, IL-33, another important IL-1 family alarmin in mucosal immunity (35), was produced in all conditions independent of the presence of PMNs. IL-33 specifically and in contrast to other IL-1 family cytokines, is released constitutively by epithelial cells where it is then processed extracellularly by serine proteases including elastase, which is released by PMNs (36). This processed form is thought to enhance inflammatory signaling. Notably, we observed significantly lower levels of IL-33 in STM-infected PMN-HIOs, which may be caused by PMN processing. To further define which cells within the PMN-HIOs contribute to inflammasome activation and IL-1 processing, paraffin sections of STM-infected PMN-HIOs were stained for ASC, an adaptor protein required for inflammasome assembly (Fig. 3C, 3D) (37). ASC-positive signal was not observed in HIO epithelial cells, consistent with a recent report that *Salmonella*-induced epithelial cell death occurs independently of ASC (28), but instead was associated with cells positive for vimentin, a protein expressed by PMNs and mesenchymal cells within the PMN-HIOs. ASC and Vimentin double-positive cells were primarily located within the lumen of PMN-HIOs, suggesting that these cells are PMNs. Closer examination of nuclear morphology of the ASC-positive cells by DAPI staining revealed multi-lobed nuclei, further supporting that inflammasome activation and IL-1 processing occur in PMNs. Together, these findings are consistent with a model where PMNs are the primary site of Caspase-1 dependent inflammasome activation and the production of IL-1 family cytokines during infection in the PMN-HIO model.

**Fig. 3.**
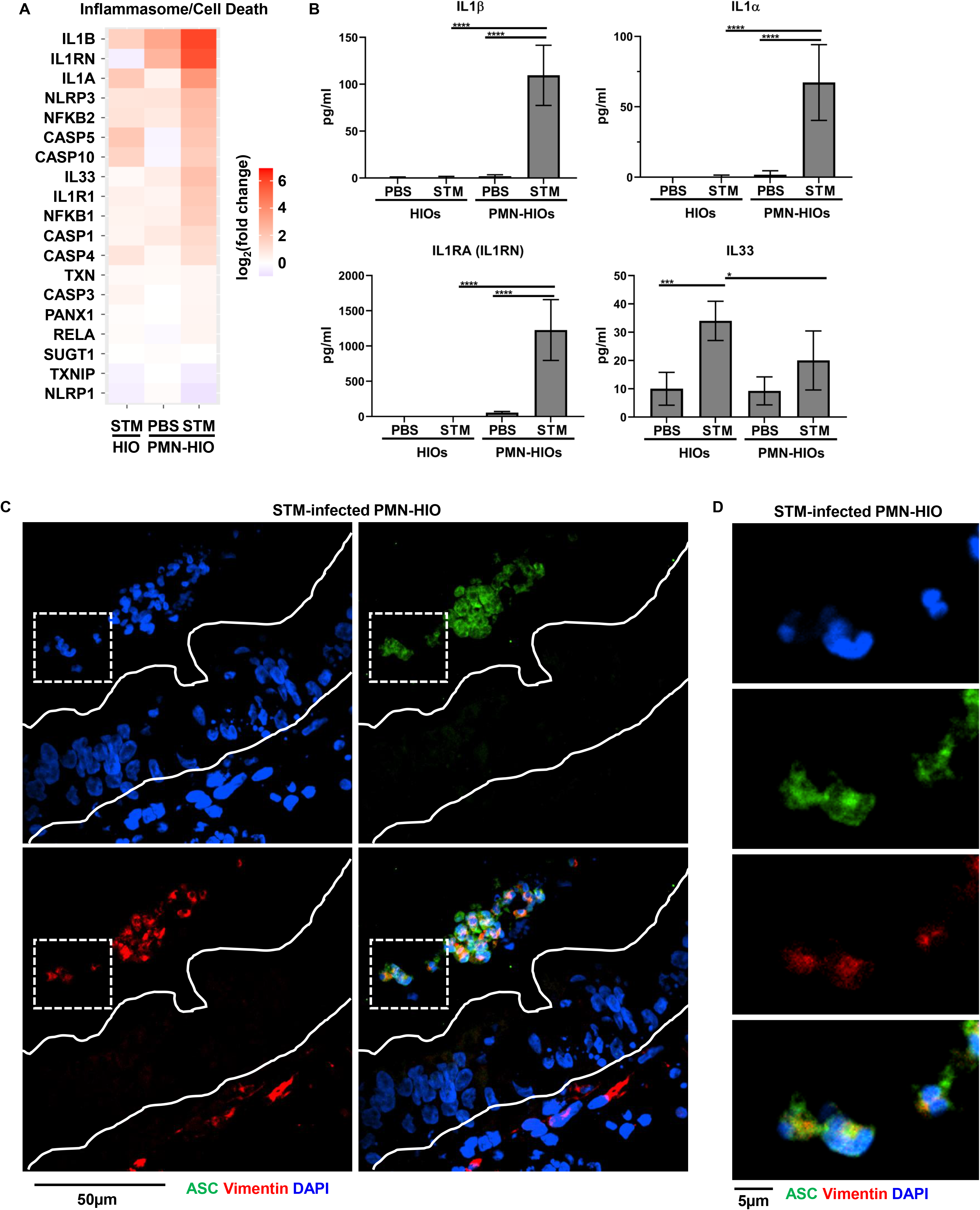
Inflammasome activation and IL-1 production is mediated by PMNs during infection. (A) Gene expression data presented as log_2_(fold change) relative to PBS-injected HIOs for genes involved in inflammasome/cell death signaling. All genes are significantly changed from PBS-injected HIOs in at least one condition with p-adjusted value <0.05. (B) Cytokine levels in culture media of HIOs and PMN-HIOs were quantified using ELISA. Graphs indicate the mean of n=4 biological replicates +/− SD from media sampled at 8hpi with 5 HIOs or PMN-HIOs per well. (C) Immunofluorescent staining of histology sections of PMN-HIOs. Sections were stained for ASC expression (green), Vimentin (red) to mark PMNs and mesenchymal cells, and DNA (blue) was labeled with DAPI. (D) Zoom of (C) showing luminal ASC-positive cells (green) with multilobed PMN nuclei. Statistical significance was determined by 2-way ANOVA where *p<0.05, ***p<0.001, ****p<0.0001.

### Caspase-1 and Caspase-3 inhibition reduces shedding of epithelial cells and increases the association of *Salmonella* with the epithelium

Epithelial cell death and shedding serve to reduce bacterial burden in the intestinal epithelium (5, 6, 8–11). To define the functional consequences of PMN-induced epithelial cell death on host defense and determine which caspases were involved, we treated PMN-HIOs with various caspase inhibitors. Accumulation of TUNEL-positive epithelial cells was monitored in infected PMN-HIOs in the presence or absence of Caspase-1 (z-YVAD-FMK) or Caspase-3 (z-DEVD-FMK) inhibitors. We performed TUNEL staining on paraffin sections from infected PMN-HIOs at 8hpi (Fig. 4A, 4B). Both Caspase-1 and Caspase-3 inhibition significantly reduced accumulation of TUNEL-positive cells in the lumen of infected PMN-HIOs, indicating that PMN-dependent Caspase-1 and −3 activation is required for efficient shedding of epithelial cells. To test how caspase inhibition and therefore reduced shedding affected STM infection, PMN-HIOs were stained with an anti-*Salmonella* antibody to characterize the localization of bacteria in the HIO by quantifying the percentage of infected cells, number of bacteria per cell, and epithelium associated bacteria (Fig. 4C–E, SI Appendix, Fig. S5). Consistent with our hypothesis that PMNs enhance shedding of epithelial cells through caspase activation, there were greater numbers of bacteria per cell in Caspase-1 inhibitor-treated PMN-HIOs, but surprisingly not in Caspase-3 inhibitor-treated PMN-HIOs (Fig. 4D). Caspase-4/5 activation in human epithelial cells is known to be important in shedding of *Salmonella*-infected cells and suggests that PMNs may enhance epithelial Caspase-4/5 signaling via Caspase-1 activity to enhance cell shedding instead of utilizing Caspase-3 for this process. We also observed a trending increase in the percentage of infected cells with Caspase-1 inhibition (SI Appendix, Fig. S5). In contrast, when PMN-HIOs were treated with the Caspase-3 inhibitor, there was a significant increase in epithelium associated bacteria (Fig. 4D). There is some evidence that activated PMNs induce apoptosis of intestinal epithelial cells (27) and so NET formation during *Salmonella* infection in PMN-HIOs may result in enhanced epithelial cell shedding independent of PMN Caspase-1 to increase the rate of cell turnover and therefore reduce the association of bacteria with the apical surface of the epithelium which would protect the epithelium from future bacterial invasion. These data suggest that both Caspase-1 and Caspase-3 inhibition reduce accumulation of dead cells in the PMN-HIO lumen, but Caspase-1 activity is important for directly regulating epithelial bacterial burden while Caspase-3 reduces association of bacteria with the epithelium to protect the epithelium from the next round of infection. Taken together, our data support a model where Caspase-1 dependent inflammasome activation in PMNs enhances epithelial cell shedding via two independent pathways to control *Salmonella* infection.

**Fig. 4.**
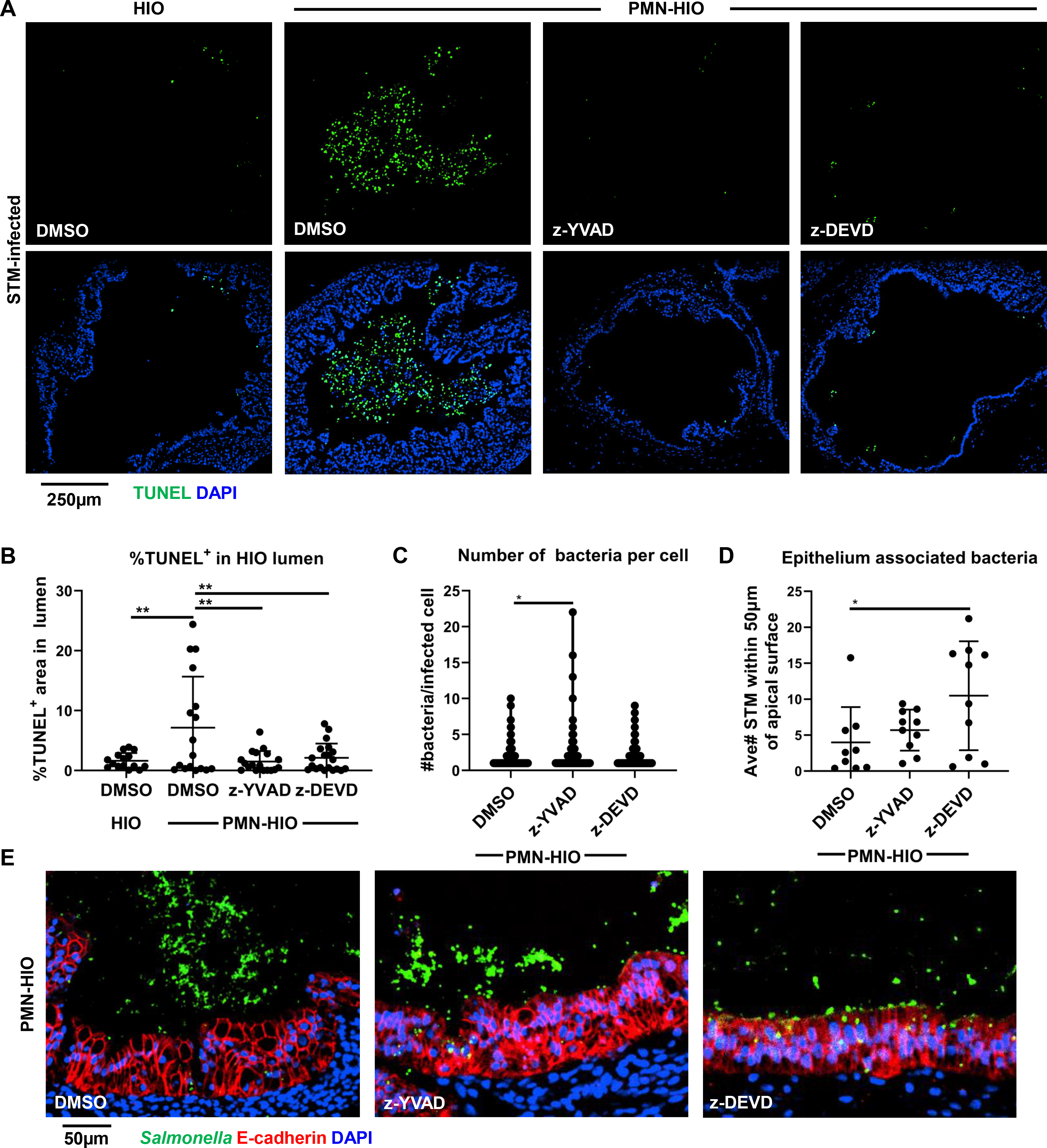
Caspase-1 and Caspase-3 inhibition reduces shedding of infected epithelial cells in the lumen of PMN-HIOs and differentially affect bacterial burden and bacterial association with the epithelium. (A) Representative fluorescence microscopy images of TUNEL staining of HIO and PMN-HIO histology sections. HIOs were microinjected with STM and either cultured alone or co-cultured with PMNs in the presence of inhibitors for Caspase-1 (z-YVAD), Caspase-3 (z-DEVD), or DMSO control. (B) Quantitation of the percent of lumen filled with TUNEL-positive cells of STM-infected HIOs or PMN-HIOs with indicated treatments. (C) Quantitation of the number of bacteria per cell based on 3 fields per view per HIO. (D) Quantitation of epithelium associated bacteria. Number of bacteria within 50μm of the apical epithelial surface were counted and normalized per 100μm distance. (E) Fluorescent microscopy images of STM-infected PMN-HIO histology sections. Samples were stained for *Salmonella* (green), E-cadherin (red), and DAPI (blue). Unless otherwise stated, graphs show the mean +/-SD of n≥ 10 HIOs represented by dots from at least two independent experiments. Outliers were removed using the ROUT method with Q=0.1%. Significance was determined by one-way ANOVA with post-Tukey’s test for multiple comparisons where *p<0.05, **p<0.01.

## Discussion

Neutrophils (PMNs) dominate the early response to *Salmonella* infection in the gut (15, 38–40), but their functions in regulating intestinal epithelial cell host defense and infection outcome are not well understood. Here we used a co-culture model of human intestinal organoids (HIOs) with primary human PMNs, termed PMN-HIOs, to elucidate these roles. We found that while there was no difference in luminal colonization of *Salmonella* in the HIOs, PMNs did reduce intraepithelial bacterial burden in the epithelium and reduced the association of *Salmonella* with the apical surface of the epithelial monolayer. PMNs were associated with elevated epithelial cell death and promoted extrusion of these cells into the lumen of infected PMN-HIOs. We found that Caspase-1 activation was required for epithelial shedding, noting that the ASC inflammasome adaptor was only present in PMNs, and that inhibition of Caspase-1 activity increased bacterial burden in HIO epithelial cells. Independently, we uncovered an important role for Caspase-3 activation in epithelial cells in reducing association of *Salmonella* with the apical surface, which was enhanced by the presence of neutrophils. Thus, we propose a model where PMNs enhance shedding of epithelial cells from the intestinal barrier via two distinct mechanisms to reduce intracellular bacterial loads and adherence of *Salmonella* to the epithelium, potentially tilting infection outcome favorably toward the host.

Our observation that PMNs did not affect total bacterial colonization in the HIO lumen was rather unexpected, both because antimicrobial effectors were produced in the PMN-HIO cultures and our findings, and that of others, indicate that PMNs kill *Salmonella* in the absence of HIOs (17). We also observed transmigration and NET formation by PMNs in the HIO lumen, so the lack of *Salmonella* killing in the HIO lumen could not be explained by lack of PMNs at the site of infection. These observations suggest the possibility that *Salmonella* may employ specific mechanisms to overcome PMN effector functions in the HIO luminal environment. Alternatively, while PMNs can be potent killers of invading bacteria, mechanisms used by PMNs to kill pathogens are not selective and therefore PMN activation near the epithelial barrier may be tightly regulated to avoid tissue damage (41). Consistent with this idea, we observed robust production of Elafin (Fig. S3), which is annotated as an antimicrobial peptide in Reactome (42). Neutrophil elastase strongly contributes to the antimicrobial function of PMNs for both intracellular killing via phagocytosis and decorating NETs (17, 43, 44). Elafin can also inhibit neutrophil elastase to reduce tissue damage caused by neutrophil overactivation (45, 46). Thus, expression of host Elafin in the presence of PMNs may trade off the ability to kill luminal *Salmonella* with protection of epithelial barrier integrity.

Addition of PMNs to the infected HIOs resulted in decreased intracellular bacterial burden, which was unexpected since intracellular bacteria are often considered protected from neutrophil killing. We therefore hypothesized that this result was due to extrusion of infected cells from the epithelial monolayer, a process mediated by caspase activation. Previous reports have highlighted the importance of epithelial cell extrusion in preventing dissemination of *Salmonella* beyond the intestine and that there are epithelial cell-intrinsic mechanisms to rid the epithelial lining of *Salmonella* (5, 6, 9–11, 47). Of note, the role of the inflammatory caspases in these processes likely differs between human and mouse, with caspase-1 playing a prominent role in mouse epithelium, while Caspase-4 plays a more prominent role in human epithelium in the absence of neutrophils. Although we could detect low levels of shed epithelial cells in STM-infected HIOs alone consistent with prior studies, this phenotype was markedly enhanced in the presence of PMNs, and inhibition with a selective Caspase-1 peptide inhibitor decreased TUNEL+ cells, as well as number of bacteria per epithelial cell. These data suggest a previously unappreciated role for PMNs in enhancing cell death in *Salmonella*-infected epithelial cells and implicate Caspase-1 in that process.

In STM-infected HIOs, we observed close association of clusters of bacteria with the epithelial surface, which would presumably advantage bacterial pathogens by spatial proximity to the monolayer. Strikingly, the introduction of PMNs into the infected HIO led to dispersal of the bacterial clusters into the HIO lumen, concomitant with a substantial increase in TUNEL+ cells. There are numerous mechanisms by which PMNs can drive epithelial cell death including oxidant production, which can activate apoptotic pathways in the epithelium (48, 49). NET formation, which we observed in our infected PMN-HIOs, also can induce epithelial cell death (27). This process was reported to be largely dependent on extracellular histones released during NET formation. We therefore consider that *Salmonella*-induced NET formation in PMN-HIOs may contribute to overall shedding of epithelial cells and the dispersal of epithelial-associated bacteria. These possibilities are consistent with our findings that both infected and uninfected epithelial cells are shed from the intestinal monolayer in the infected PMN-HIOs, but not the infected HIOs alone. By increasing epithelial cell shedding, this neutrophil-dependent process contributes to reducing bacterial burden within and associated with the epithelium.

Our findings pointed to Caspase-1 activity as a driver of accumulation of shed epithelial cells in the HIO lumen, but markers of inflammasome activation were only observed when PMNs were present. Although the importance of Caspase-1 in PMN activation and antimicrobial functions are not fully understood, some evidence implicates Caspase-1 in NET formation (29, 50). Our prior studies demonstrated that inflammasome activation in PMNs occurs during bacterial infection (29) and another study showed that inflammasome-mediated processing of Gasdermin D was required for NET formation in mice (50). While there may be additional signaling roles of Caspase-1 in PMN activation, we propose a model whereby PMNs transmigrate into the inflamed intestine, undergo Caspase-1 mediated NET formation to trigger epithelial cell death and shedding of infected cells to protect the epithelium from ongoing infection. We reason that in the environment of the infected gut, in contrast to infected tissue, the ratio of commensal and pathogenic bacteria to neutrophils may preclude substantive bacterial killing through direct anti-microbial mechanisms. Therefore, signaling mechanisms whereby neutrophils are able to direct protective epithelial responses may more be more advantageous to the host.

## Materials and Methods

### Contact for Reagent and Resource Sharing

Reagents and resources can be obtained by directing requests to the corresponding authors, Basel Abuaita and Mary O’Riordan.

### Human Intestinal Organoids (HIOs)

HIOs were generated by the *In Vivo* Animal and Human Studies Core at the University of Michigan Center for Gastrointestinal Research as previously described (51). Prior to experiments, HIOs were removed from the Matrigel, washed with DMEM:F12 media, and re-plated with 5 HIOs/well in 50μl of Matrigel (Corning) in ENR media ((DMEM:F12, 1X B27 supplement, 2mM L-glutamine, 100ng/ml EGF, 100ng/ml Noggin, 500ng/ml Rspondin1, and 15mM HEPES). Media was exchanged every 2-3 days for 7 days.

### Human Polymorphonuclear Leukocytes (PMNs)

PMNs were isolated from blood of healthy human volunteers as previously described (29). The purity of PMNs was assessed by flow cytometry using APC anti-CD16 and FITC anti-CD15 antibodies (Miltenyi Biotec); markers characteristic of human neutrophils. PMNs were labeled with cell trace CFSE dye (Thermo Fisher). PMNs were incubated at room temperature for 20 minutes in PBS containing 5 μM CFSE. Cells were washed twice with PBS to remove excess dye and collected by centrifugation. CFSE-labeled PMNs were then co-cultured with STM-infected HIOs or PBS control to monitor the association of PMNs with intestinal epithelial cells. PMN-HIOs were washed twice with PBS to remove unassociated PMNs, mechanically dissociated into single-cell suspension using a 70 μm cell strainer and analyzed on FACSCanto flow cytometer. Percent of CFSE-positive cells were determined using FlowJo software.

### Bacterial Growth and HIO Microinjection

*Salmonella enterica* serovar Typhimurium SL1344 (STM) was used throughout the manuscript. Bacteria were stored at −80°C in Luria-Bertani (LB, Fisher) medium containing 20% glycerol and cultured on LB agar plates. Individual colonies were grown overnight at 37ºC under static conditions in LB liquid broth. Bacteria were pelleted, washed and re-suspended in PBS. Bacterial inoculum was estimated based on OD_600_ and verified by plating serial dilutions on agar plates to determine colony forming units (CFU). The lumen of individual HIOs were microinjected with glass caliber needles with 1μl of PBS or STM (10^5^ CFU/HIO) as previously described (3, 4, 52). HIOs were then washed with PBS and incubated for 2h at 37°C in ENR media. HIOs were treated with 100μg/ml gentamicin for 15 min to kill any bacteria outside the HIOs, then incubated in fresh medium +/− PMNs (5 × 10^5^ PMNs/5HIOs/well in a 24-well plate). Where indicated, PMNs-HIOs were treated with the following inhibitors after microinjection: 4 μM of Caspase-1 inhibitor, Z-YVAD-FMK or 4 μM Caspase-3 inhibitor, Z-DEVD-FMK.

### Bacterial Burden and Cytokine Analyses

Bacterial burden was assessed per HIO. Individual HIOs were removed from Matrigel, washed with PBS and homogenized in PBS. Total CFU/HIO were enumerated by serial dilution and plating on LB agar. For cytokine analysis, media from each well containing 5 HIOs/well were collected at 8hpi. Cytokines, chemokines and antimicrobial proteins were quantified by ELISA at the University of Michigan Cancer Center Immunology Core.

### Immunofluorescence Staining and Microscopy

HIOs were fixed with 10% neutral formalin for 2 days and embedded in paraffin. Histology sections (5μm) were collected by the University of Michigan Cancer Center Histology Core. Sections were deparaffinized and antigen retrieval was performed in sodium citrate buffer (10mM sodium citrate, 0.05% Tween 20, pH 6.0). Sections were permeabilized with PBS+ 0.2% Triton X-100 for 30 min, then incubated in blocking buffer (PBS, 5% BSA, and 10% normal goat serum) for 1h. Primary antibodies; anti-E-Cadherin (BD Biosciences, clone 36), anti-MPO (Agilent, clone A0398), anti-Vimentin (DSHB, Cat# AMF-17b), anti-ASC (Cell Signaling, Cat#13833) and anti-cleaved Caspase-3 (Cell Signaling, Cat# 9661) were added to the histology sections in blocking buffer overnight at 4°C. Goat anti-mouse and anti-rabbit secondary antibodies conjugated to Alexa-488, Alexa-594 or Alexa-647 were used according to manufacturer’s instructions (Thermo Fisher) for 1h RT in blocking buffer. DAPI (Thermo Fisher) was used to stain DNA. Bacteria were stained using anti-*Salmonella* Typhimurium FITC-conjugated antibody (Santa Cruz, Cat# sc-52223). Sections were mounted using coverslips (#1.5) and Prolong Diamond or Prolong Glass Antifade Mountant (Thermo Fisher). Images were taken on Olympus BX60 upright compound microscope, Nikon A1 confocal microscope or Nikon X1 Yokogawa spinning disc confocal microscope and processed using ImageJ and quantitation was performed in ImageJ or CellProfiler.

### TUNEL Assay

Apoptosis was analyzed by fluorescence microscopy using *In Situ Cell Death Detection Kit* (Roche) or CF594 TUNEL Assay Apoptosis Detection Kit (Biotium) according to the manufacturers’ protocols. Histology sections were permeabilized using Proteinase K (20μg/ml) or 0.2% Triton X-100 in PBS and blocked using PBS+ 5% BSA. Sections were stained with primary antibodies overnight at 4°C in blocking buffer and then were incubated in the terminal deoxynucleotidyl transferase end labeling (TUNEL) buffer for 1h at 37°C. Slides were washed with PBS and incubated with fluorescent conjugated secondary antibodies. Sections were then counterstained with DAPI to label DNA. To quantify the TUNEL signal in the HIOs, the percent of the HIO lumen filled with TUNEL+ cells was quantified using ImageJ software.

### RNA Sequencing and Analysis

Total RNA was isolated from 5 HIOs per group with a total of 4 replicates per condition using the mirVana miRNA Isolation Kit (Thermo Fisher). The quality of RNA was confirmed, ensuring the RNA integrity number (RIN)> 8.5, using the Agilent TapeStation system. cDNA libraries were prepared by the University of Michigan DNA Sequencing Core using the TruSeq Stranded mRNA Kit (Illumina) according to the manufacturer’s protocol. Libraries were sequenced on Illumina HiSeq 2500 platforms (single-end, 50 bp read length). All samples were sequenced at a depth of 10.5 million reads per sample or greater. Sequencing generated FASTQ files of transcript reads that were pseudoaligned to the human genome (GRCh38.p12) using kallisto software (53). Transcripts were converted to estimated gene counts using the tximport package (54) with gene annotation from Ensembl (55).

### Gene Expression and Pathway Enrichment Analysis

Differential expression analysis was performed using the DESeq2 package (56) with *P* values calculated by the Wald test and adjusted *P* values calculated using the Benjamani & Hochberg method (57).

### Quantification and Statistical Methods

RNA-seq data analysis was done using RStudio version 1.1.453. Plots were generated using ggplot2 (58) with data manipulation done using dplyr (59). Other data were analyzed using Graphpad Prism 9. Statistical differences were determined using statistical tests indicated in the fig. legends. The mean of at least 2 independent experiments were presented with error bars showing standard deviation (SD). *P* values of less than 0.05 were considered significant and designated by: \**P* < 0.05, \*\**P* < 0.01, \*\*\**P* < 0.001 and **** *P* < 0.0001.

### Ethics Statement

Blood samples were obtained from healthy adult donors according to the protocol approved by the University of Michigan Institutional Review Board (HUM00044257). Written consent was obtained from all donors.

## Acknowledgements

This work was supported by NIH awards U19AI116482-01 (V.B.Y, J.R.S, C.E.W, and M.X.O), R21AI13540 (M.X.O) and funds from the University of Michigan Medical School (J.S.K). A-L.E.L was supported by NIH T32 AI007528. We thank the Host Microbiome Initiative, Microscopy and Image Analysis Laboratory (MIL), the Comprehensive Cancer Center Immunology (supported by the NCI and NIH award P30CA046592) and Histology Cores and the DNA Sequencing Core at University of Michigan Medical School. We gratefully acknowledge the O’Riordan lab members for helpful discussions, and Dr. Roberto Cieza for assistance with data management.

---

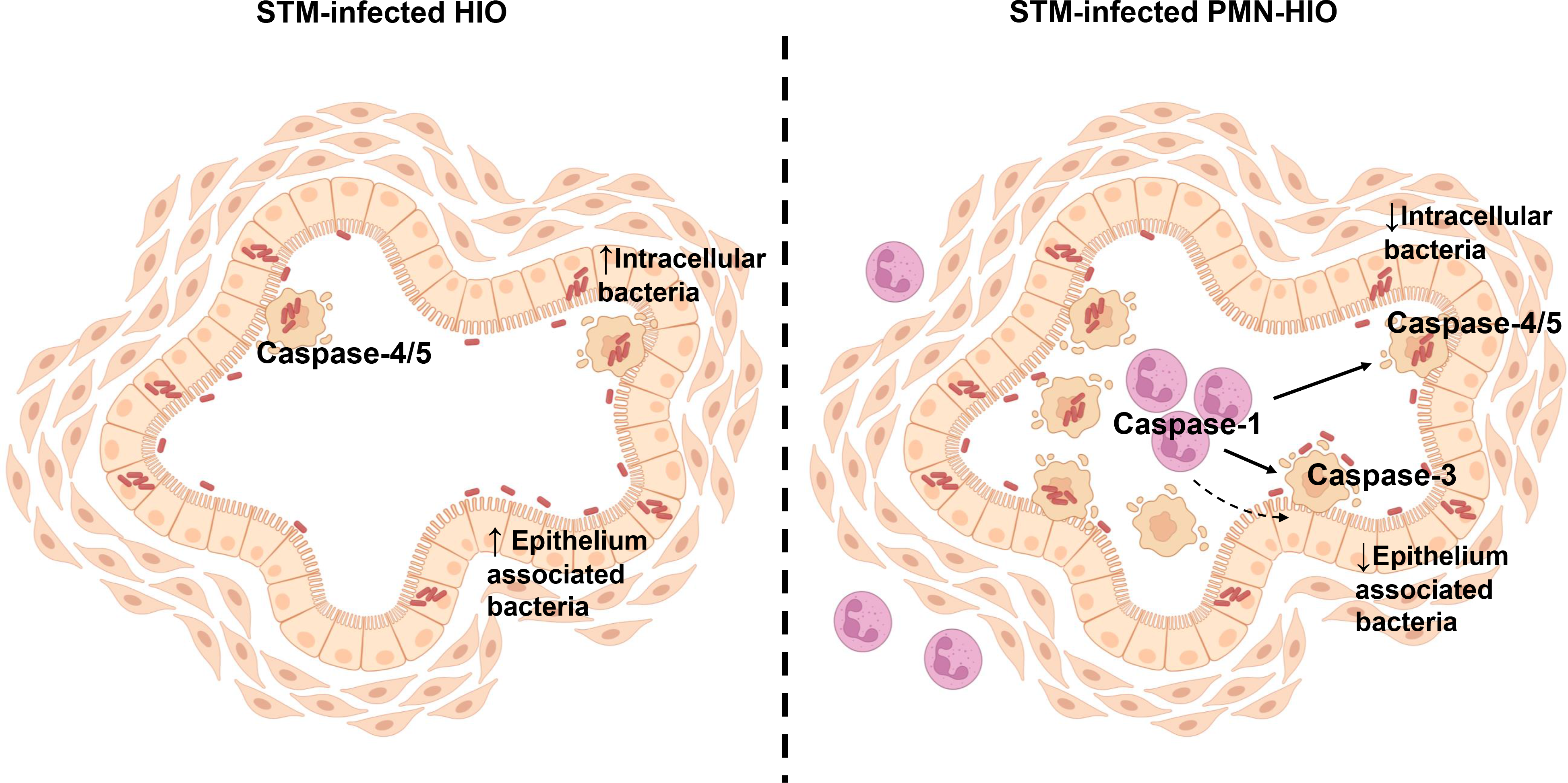

## Notes

### Competing Interest Statement

The authors have declared no competing interest.

### Summary of Updates

Supplemental Material added.

