## Supplemental Information for "Human neutrophils direct epithelial cell extrusion to enhance intestinal epithelial host defense during *Salmonella* infection"

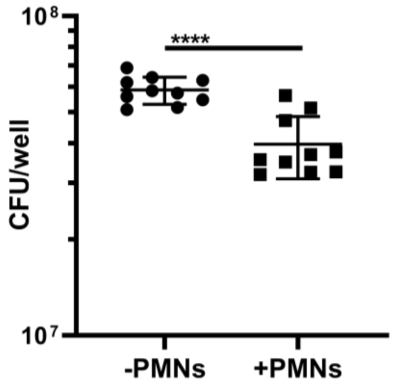


**Figure S1. PMNs kill STM**

PMN bactericidal activity against *Salmonella* was quantified by enumerating CFU at 4h in the presence of PMNs relative to bacteria cultured alone. Results are from n=4 independent experiments with PMNs isolated from blood of different donors.

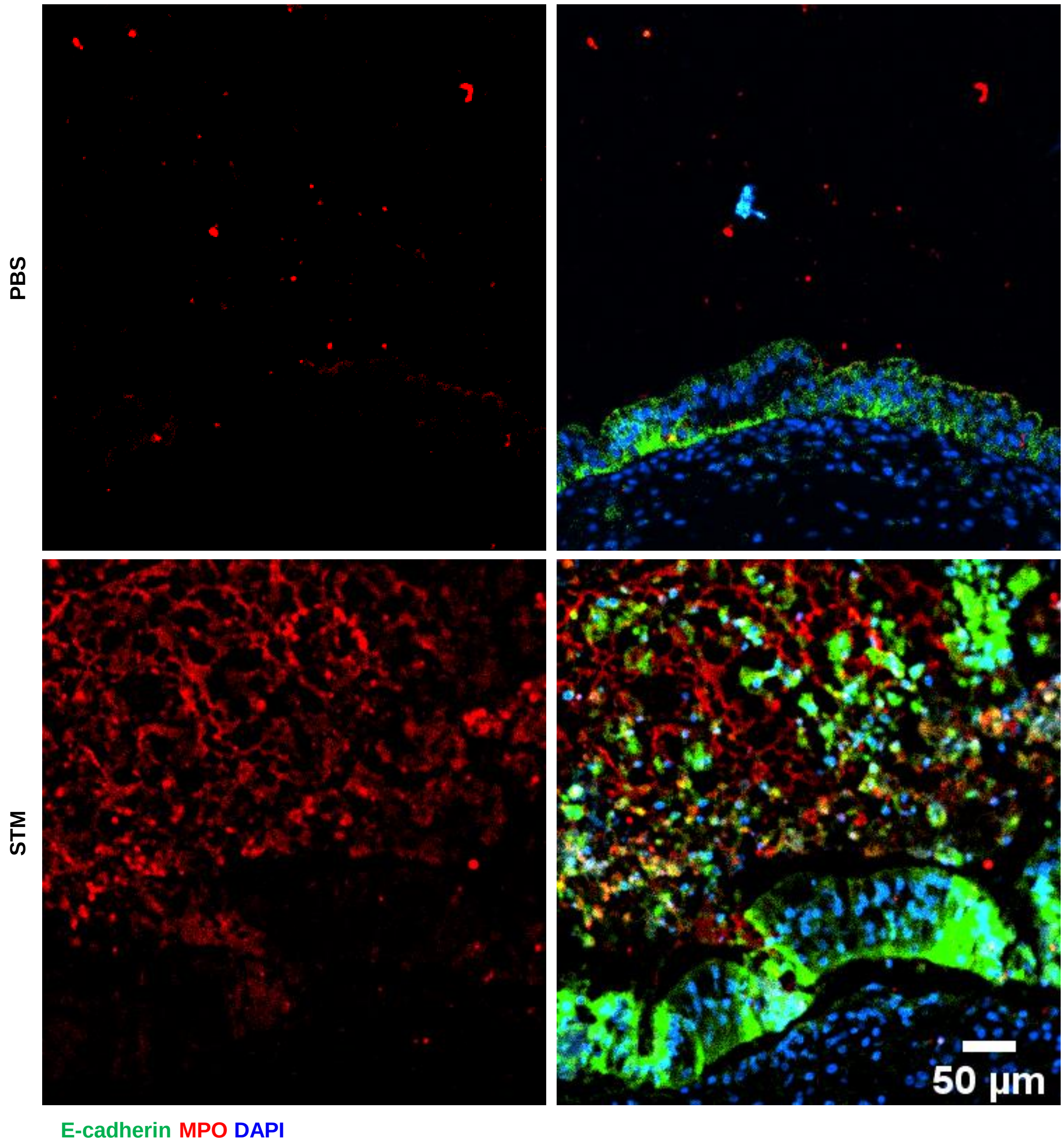


**Figure S2. PMNs form NETs in PMN-HIOs during *Salmonella* infection**
Immunofluorescent staining of PMN-HIOs microinjected with PBS or STM and stained for epithelial cells marked with E-cadherin (green), PMNs marked by MPO (red) and DNA with DAPI (blue).

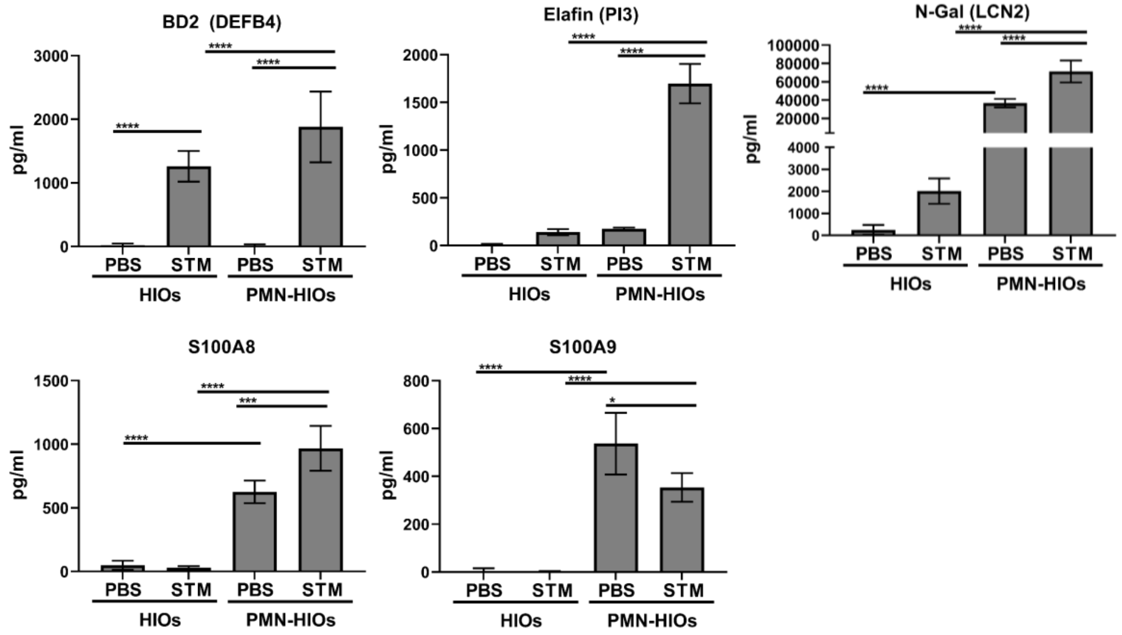


**Figure S3. The antimicrobial response is intact in PMN-HIOs**
Quantitation of antimicrobial protein levels in culture media of HIOs and PMN-HIOs microinjected with PBS or STM for 8h measured by ELISA. Graphs indicate the mean of n=4 replicates +/- standard deviation. Significance was determined by 2-way ANOVA where *p<0.05, ***p<0.001, ****p<0.0001.

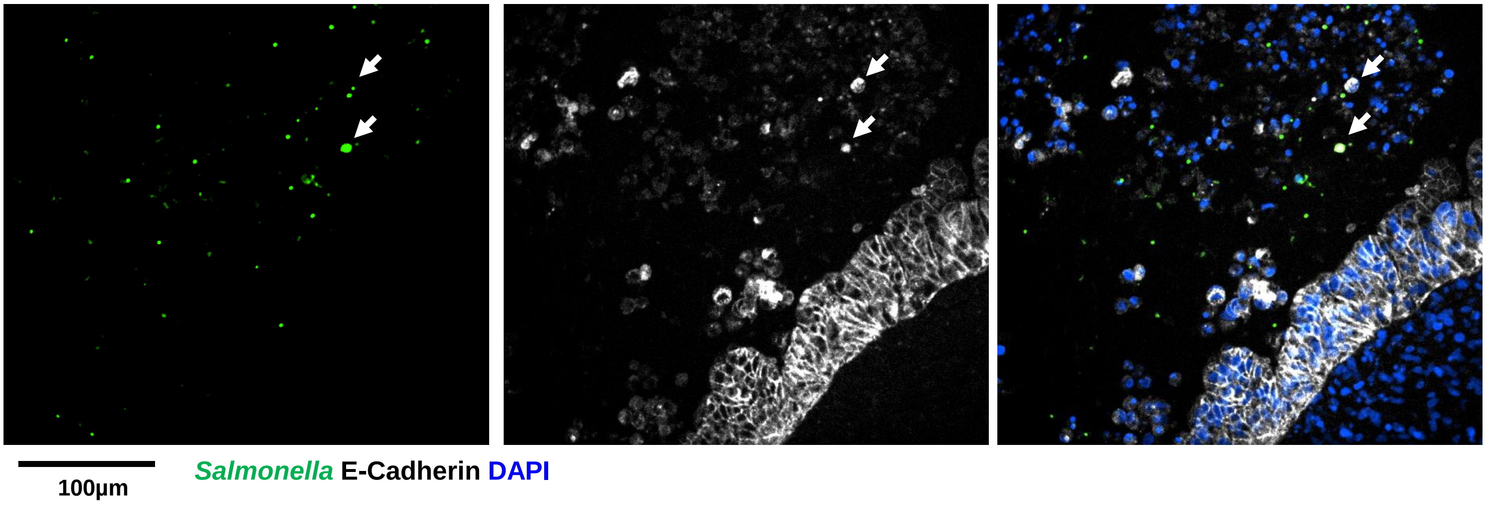


**FigureS4. Some but not all extruded cells are infected with *Salmonella***
Immunofluorescent staining of STM-infected PMN-HIO stained for *Salmonella* (green), E-cadherin (white), and DAPI (blue). Arrowheads point to infected extruded cells.

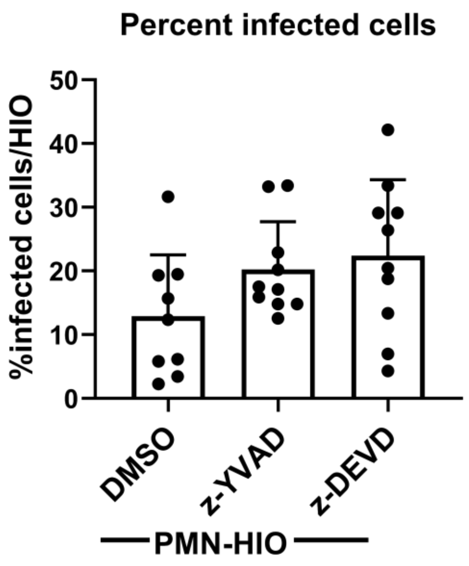


**Figure S5 Percent infected cells with Caspase-1 or Caspase-3 inhibition**
Quantitation of percent infected cells/HIO or PMN-HIO based on 3 fields per view per HIO. Graphs show the mean and SD of n≥ 10 HIOs represented by dots from at least two independent experiments.
